# Epi-mutations for spermatogenic defects by maternal exposure to Di (2-ethylhexyl) phthalate

**DOI:** 10.1101/2021.05.19.444770

**Authors:** Yukiko Tando, Hitoshi Hiura, Asuka Takehara, Yumi Ito-Matsuoka, Takahiro Arima, Yasuhisa Matsui

## Abstract

Exposure to environmental factors during fetal development may lead to epigenomic modifications in fetal germ cells, altering gene expression and promoting diseases in successive generations. In mouse, maternal exposure to Di (2-ethylhexyl) phthalate (DEHP) is known to induce defects in spermatogenesis in successive generations, but the mechanism(s) of impaired spermatogenesis are unclear. Here, we showed that maternal DEHP exposure results in DNA hypermethylation of promoters of spermatogenesis-related genes in fetal testicular germ cells in F1 mice, and hypermethylation of *Hist1h2ba, Sycp1* and *Taf7l,* which are crucial for spermatogenesis, persisted from fetal testicular cells to adult spermatogonia, resulting in the downregulation of expression of these genes. Forced methylation of these gene promoters silenced expression of these loci in a reporter assay. Expression and methylation of those genes tended to be downregulated and increased, respectively in F2 spermatogonia following maternal DEHP exposure. These results suggested that DEHP-induced hypermethylation of *Hist1h2ba, Sycp1* and *Taf7l* in fetal germ cells results in downregulation of these genes in spermatogonia and subsequent defects in spermatogenesis, at least in the F1 generation.

## Introduction

A wide variety of environmental influences have been shown to greatly impact human health, with the effects persisting through multiple generations (Daxinger & Whitelaw, 2012; Taouk & Schulkin, 2016). Historically, studies on the Dutch famine of 1944 indicated that poor maternal nutrition during pregnancy was associated with low birth weight as well as a greater risk of metabolic and cardiovascular diseases in offspring (Painter *et al*., 2005). Those studies also showed that maternal nutritional conditions resulted in changes in offspring in DNA methylation patterns in the promoters of genes related to metabolic diseases and cardiac diseases (Tobi *et al*., 2009), suggesting the involvement of epigenetic modifications in germ cells in inter-/trans-generational influences of the maternal environment. In a rodent model, a maternal nutritional change induced increased DNA methylation of CpG sites in the *A^vy^* allele and inhibition of expression of the corresponding gene, which in turn may result in changes in the hair color of offspring (Waterland & Jirtle, 2003).

With regard to paternal influences, various environmental factors also affect traits of offspring in human as well as rodents (Pembrey *et al*., 2006; Ng *et al*., 2010). For instance, studies of paternal inflammatory and nutritional modifications suggested that the environmentally induced epigenetic alterations in sperm can be inherited by the next generation. Specifically, chemically induced liver damage has been shown to influence paternally inherited suppressive adaptation for liver fibrosis in F1 and F2 generations; these effects are mediated by epigenetic changes in sperm, and up- and down-regulation of anti- and pro-fibrogenic genes, respectively, in offspring (Zeybel *et al*., 2012). Concerning nutritional influences, paternal prediabetes has been shown to alter DNA methylation of sperm genes encoding components of the insulin signaling pathway, leading to glucose intolerance and insulin resistance in the offspring (Wei *et al*., 2014). Another example is that offspring of males fed a low-protein diet (LPD) exhibit upregulation of genes involved in lipid and cholesterol biosynthesis and decreased levels of cholesterol esters in the liver; these offspring also exhibit changes in the DNA methylation of genes involved in the regulation of lipid metabolism (Carone *et al*., 2010). In addition, LPD-induced epigenetic changes (such as histone H3 lysine (K) 9 methylation levels and small RNA expression) in testicular germ cells have been shown to be mediated by cyclic AMP-dependent transcription factor 7 (ATF7), with resulting effects on gene expression in offspring (Yoshida *et al*., 2020).

As in mammals, transgenerational inheritance of the effects of environmental factors via epigenetic modifications has been demonstrated in model organisms such as *Drosophila* and *Caenorhabditis elegans.* Heat shock or osmotic stress in adult or embryonic *Drosophila* induces phosphorylation of Drosophila activating transcriptional factor-2 (dATF-2) and results in the release of this protein from heterochromatin; this heterochromatic disruption is an epigenetic event that is transmitted to the next generation in a non-Mendelian fashion (Seong *et al*., 2011). In *C. elegans,* starvation-induced developmental arrest leads to production of small RNAs that are inherited through at least three consecutive generations; these small RNAs target genes with roles in nutrition (Rechavi *et al*., 2014).

Transgenerational defects in spermatogenesis has been demonstrated after maternal exposure to several chemicals, including vinclozolin (Anway *et al*., 2005); arsenic (Xia *et al*., 2021); and p,p’-dichlorodiphenoxydichloroethylene (p,p’-DDE), one of the primary metabolic products of the classical organochlorine pesticide, dichlorodiphenoxytrichloroethane (DDT) (Song *et al*., 2014). Recent work has demonstrated changes of DNA methylation in fetal germ cells and spermatogenetic cells in postnatal testis after prenatal exposure to vinclozolin (Skinner *et al*., 2019) and DDT (ben Maamar *et al*., 2019), but a causal relationship between the altered DNA methylation/gene expression and the spermatogenetic defects has not been determined.

Di (2-ethylhexyl) phthalate (DEHP) is a plasticizer that is used in a wide range of consumer products, such as food packing, medical devices, and wallpapers. Maternal exposure to DEHP has been shown to affect spermatogenesis in male offspring (Doyle *et al*., 2013) as well as several other body systems over multiple generations, including the female reproductive system (Brehm *et al*., 2018), liver (Wen *et al*., 2020), and anxiety-like behavior (Quinnies *et al*., 2015). Concerning influences on spermatogenic cells, offspring that are exposed prenatally to DEHP show defects in spermatogenesis, decreases in sperm number, and reduction of sperm motility (Doyle *et al*., 2013; Prados *et al*., 2015; Stenz *et al*., 2017), and F1 sperm demonstrate altered DNA methylation and expression of the genes encoding seminal vesicle secretory protein 2 (Svs2; also known as semenogelin-1); Svs3b; Svs4; and seminal vesicle antigen (Sva), all of which are involved in sperm motility (Stenz *et al*., 2017). However, possible causes of inter- or trans-generational spermatogenic defects, including epi-mutations of spermatogenesis-related genes, have not been identified; such causes are expected to be essential to understanding the mechanism(s) of the transgenerational effects of DEHP on spermatogenesis. In the present study, we examined changes in DNA methylation and gene expression in fetal male germ cells shortly after maternal DEHP exposure and in spermatogenic cells in adult offspring. We identified candidate epi-mutated genes that are involved in abnormal spermatogenesis in mouse.

## Results

### Defects in spermatogenesis following maternal DEHP exposure

Using C57BL/6 mice, we confirmed the previously reported effects of maternal DEHP exposure on testicular germ cells in embryos and in testes in subsequent generations. First, we used histological evaluation to examine spermatogenesis in the F1 of dams exposed to DEHP and in the F2 resulting from mating of these maternally exposed F1 males with untreated C57BL/6 females. Testicular abnormalities, including vacuoles in tubules (Type A), no lumen in tubules (Type B), and apoptotic cells (Type C), were observed in F1 males of the DEHP group, while few abnormal tubules were found in the oil group as vehicle control (Fig 1A), as previously reported (Doyle *et al*., 2013). In Type-A tubules, the size and number of vacuoles varied among tubules. In tubules with large vacuoles, germ cells were typically deleted. In tubules with small vacuoles-1, spermatocytes and spermatids were decreased in number, and no spermatozoa were found. In tubules with small vacuoles-2, spermatogenesis was arrested at the spermatocyte stage. In tubules with small vacuoles-3, abnormal spermatids with peripherally positioned nuclei were found (Fig 1A, enlarged images). F2 males of the DEHP group showed similar histological abnormalities (Fig 1B).

**Figure 1.**
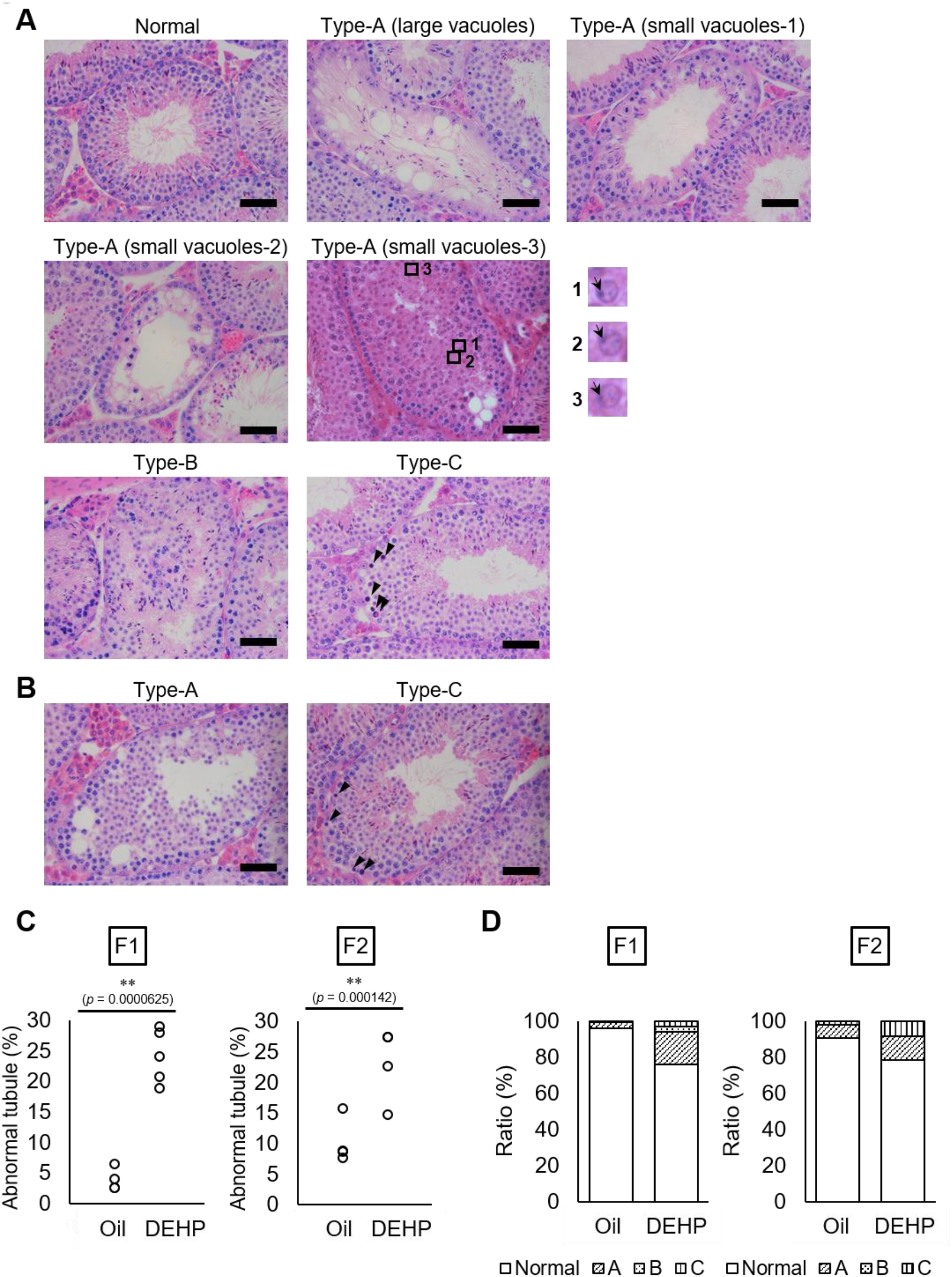
Histological analysis of testicular tubules of C57BL/6 offspring after prenatal DEHP exposure. A, B. Representative images of testicular tubules in F1 (A) and F2 (B). Abnormal testicular tubules were categorized as “Type A” (presence of various size of vacuoles), “Type B” (no lumen and complete loss of germ cell organization), or “Ty pe C” (presence of dead cells; indicated by arrowheads). Type A was further classified into three subtypes as described in Results. Enlarged images corresponding to the rectangular areas in Type A (small vacuoles-3) are shown. Arrows indicate nuclei. Arrowheads indicate apoptotic cells. Scale bars: 50 μm. C. Ratios of abnormal tubules in F1 and F2 animals. (F1 oil as vehicle control: n=4; F1 DEHP: n=6; F2 oil: n=4, F2 DEHP: n=4). ** *P* < 0.01 (unpaired two-sided Student’s t-test). Data are presented as mean ± SEM. D. Ratios of abnormal tubule types in F1 and F2.

Quantitative evaluation of the abnormal testicular tubules showed 6- and 2-fold increases of the ratios of the abnormal tubules in the DEHP-exposed F1 and F2 testes, respectively, compared with those in the oil-exposed groups (Fig 1C). Classification of the types of the abnormalities, as shown in Fig 1A, revealed that Type B was not observed in F2 (Fig 1D), suggesting some difference in the nature of spermatogenic failure between the F1 and the F2 testes. We also observed similar testicular abnormalities in F1 and F2 males derived from Oct4-deltaPE-GFP transgenic mice with a C57BL/6 background, which were used for isolating fetal testicular germ cells (figure supplement 1A-D). In addition, a significant increase in the number of multinucleated germ cells was observed in embryonic day (E) 19.5 testis in animals prenatally exposed to DEHP compared with those in an oil-exposed group (figure supplement 1E, F), as reported previously (Ungewitter *et al*., 2017). These results indicated that our maternal DEHP exposure experiments reproduced the spectrum of testicular abnormalities described in a previous report (Doyle *et al*., 2013).

### Changes of DNA methylation in fetal and adult testicular germ cell populations of F1 following maternal DEHP exposure

To determine DNA methylation changes in germ cells following maternal DEHP exposure, we performed reduced representation bisulfite sequencing (RRBS) of DNA from germ cells purified from E19.5 testes and from adult F1 testes at approximately postnatal day (P) 200. Germ cells and gonadal somatic cells in E19.5 testes were purified as Oct4-deltaPE-GFP-positive cells by flow cytometry (figure supplement 2A). Spermatogonia, spermatocytes, and round spermatids were purified from adult testes by flow cytometry based on their profiles when stained with Hoechst dye, and enrichment of each cell type was confirmed by screening for the expression of known marker genes *(Gfra1* for spermatogonia, *Scp3* for spermatocytes, and *Acrv1* for spermatids; figure supplement 2B, C). Comparison of mean methylation levels in CpG cytosines in various genomic features (including promoters, gene bodies, CpG islands, CpG island shores, and transposons) showed similar methylation levels between the oil- and the DEHP-exposed groups in each germ cell population in all genomic regions (Supplementary Table 1). Heatmap analysis also showed moderate differences in promoter methylation profiles between the oil and the DEHP groups (Fig 2A). At the same time, the methylation profiles differed strikingly between spermatogonia and other germ cell populations, such that the promoters in spermatogonial DNA tended to be hyper-methylated compared to those in other cell populations, a result that is consistent with the mean methylation levels shown in Supplementary Table 1.

**Figure 2.**
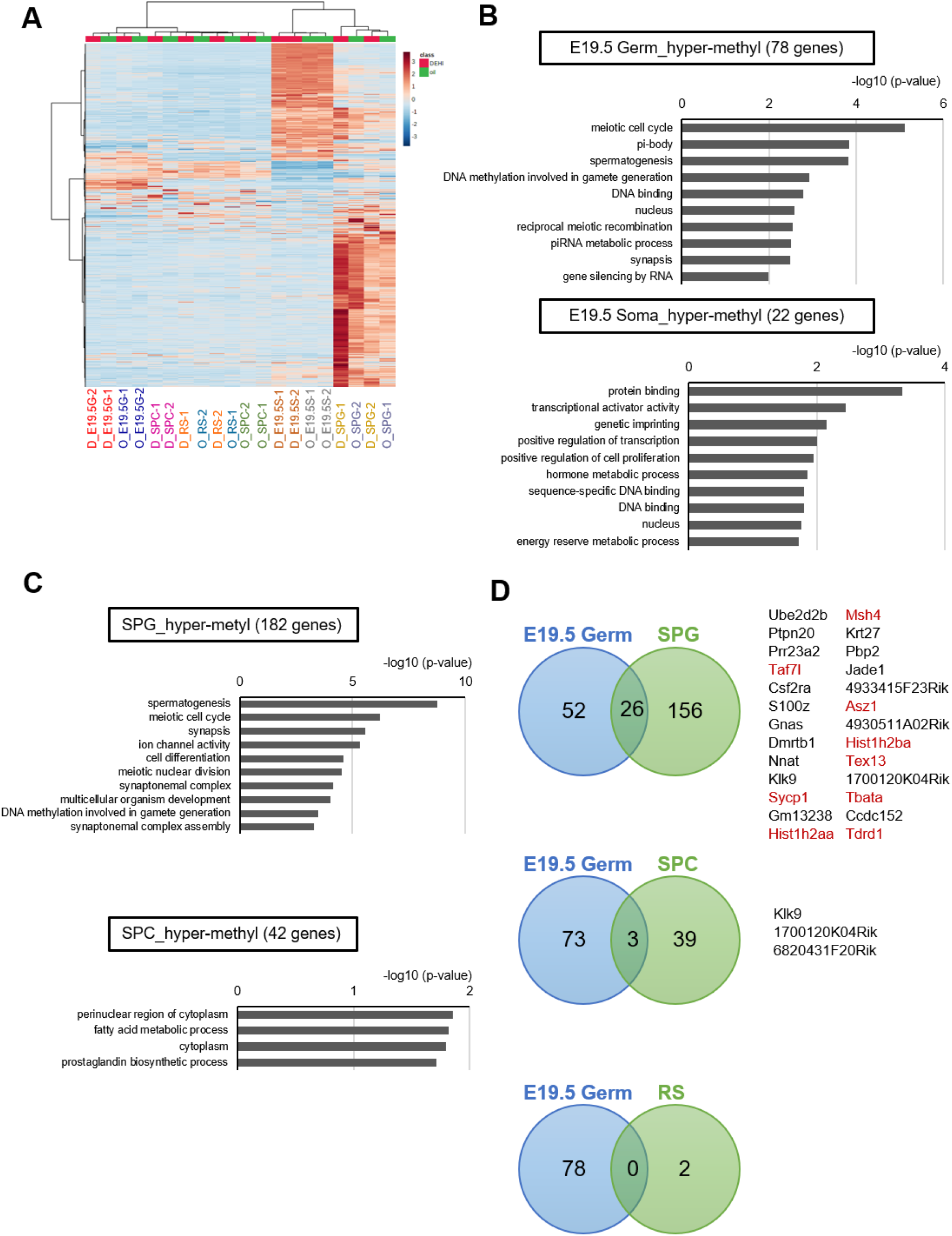
Methylome of testicular cells of F1 after maternal exposure to DEHP. A. Heatmap of promoter DNA methylation values in the testicular cell populations prenatally exposed to oil as vehicle control (O) or DEHP (D). E19.5G: E19.5 germ cell, E19.5S: E19.5 testicular somatic cells, SPG: spermatogonia; SPC: spermatocytes. B, C. Functional annotation of hypermethylated (more than 5% increase in DEHP-treated samples compared to oil-treated samples) genes in E19.5 germ cells (Germ) and testicular somatic cells (Soma) (B) and in adult germ cell populations (C) using DAVID. D. Venn diagram analysis of the hypermethylated genes in E19.5 germ cells, spermatogonia (SPG), spermatocytes (SPC), or round spermatids (RS). Genes that are hypermethylated in E19.5 germ cell and SPG or SPC are listed, and spermatogenesis-related genes are shown in red.

We next attempted to identify differentially methylated regions (DMRs) in the DNA of E19.5 germ cells when comparing between the oil and DEHP groups. This analysis identified 80 and 22 promoters (78 and 20 genes) with hyper- and hypo-methylation, respectively, in DEHP groups (compared to oil groups; Supplementary Table 2). These DMRs were widely distributed across the genome in both germ cells and testicular somatic cells (figure supplement 3). Functional enrichment analysis revealed that spermatogenesis-related GO (gene ontology) terms such as meiotic cell cycle and male gamete generation were enriched in hypermethylated genes in germ cells, while such enrichment was not observed in hypermethylated genes in testicular somatic cells (Fig 2B, Supplementary Table 2). In addition, specific functional terms were not enriched in the hypomethylated genes shown in Supplementary Table 2. Together, these results suggested preferential hypermethylation of spermatogenesis-related genes in E19.5 germ cells following maternal DEHP exposure.

We also compared methylation in F1 testicular germ cells, and identified 191, 43, and 2 promoter regions (182, 42, 2 genes) with hypermethylation in spermatogonia, spermatocytes, and round spermatids, respectively, in the DEHP group compared with those in the oil group, among which spermatogenesis-related genes were enriched only in spermatogonia (Fig 2C, Supplementary Table 3). As in E19.5 germ cells, only a small number of hypomethylated genes were identified, and functional GO terms were not enriched for these genes (Supplementary Table 3).

We then examined whether the hypermethylated promoter regions detected in the E19.5 germ cells were maintained in F1 testicular germ cells (Fig 2D). We found 26 genes that were hypermethylated in both E19.5 germ cells and F1 spermatogonia; of these 26 genes, 9 encoded products related to spermatogenesis. Meanwhile, no hypermethylated and spermatogenesis-related genes were found to be shared between the E19.5 germ cells and spermatocytes or round spermatids (Fig 2D). These results implied that the DEHP-induced hypermethylation in spermatogenesis-related genes is maintained up to the spermatogonial stage in F1 testis.

### Changes in the expression of the hyper-methylated spermatogenesis-related genes in F1 spermatogonia

We next examined changes in gene expression in E19.5 and F1 testicular germ cells following maternal DEHP exposure. Overall, gene expression profiles obtained by RNA-seq showed no remarkable differences between the oil and DEHP groups in each germ cell population (figure supplement 4A). GO analysis of the differentially expressed genes (DEGs) between the oil and the DEHP groups showed enrichment of several biological processes in E19.5 germ cells as well as in F1 testicular germ cell populations (figure supplement 4B, C, Supplementary Table 4). Enrichment of germ cell-related GO terms was observed in upregulated genes in spermatogonia and spermatocytes. It seems to be inconsistent with the enrichment of spermatogenesis-related GO terms in hypermethylated genes in spermatogonia (Fig 2C), but genes included in spermatogenesis GO for hypermethylated or upregulated genes by DEHP in spermatogonia are not overlapping (figure supplement 4D). In addition, upregulated genes in spermatogonia and spermatocytes are mainly associated with spermiogenesis, while hypermethylated genes in spermatogonia are involved in meiosis (Fig 3C, figure supplement 4C). Taken together, spermiogenesis-related genes may be abnormally upregulated in spermatogonia by DEHP exposure, which is not directly controlled by DNA methylation status of the genes.

**Figure 3.**
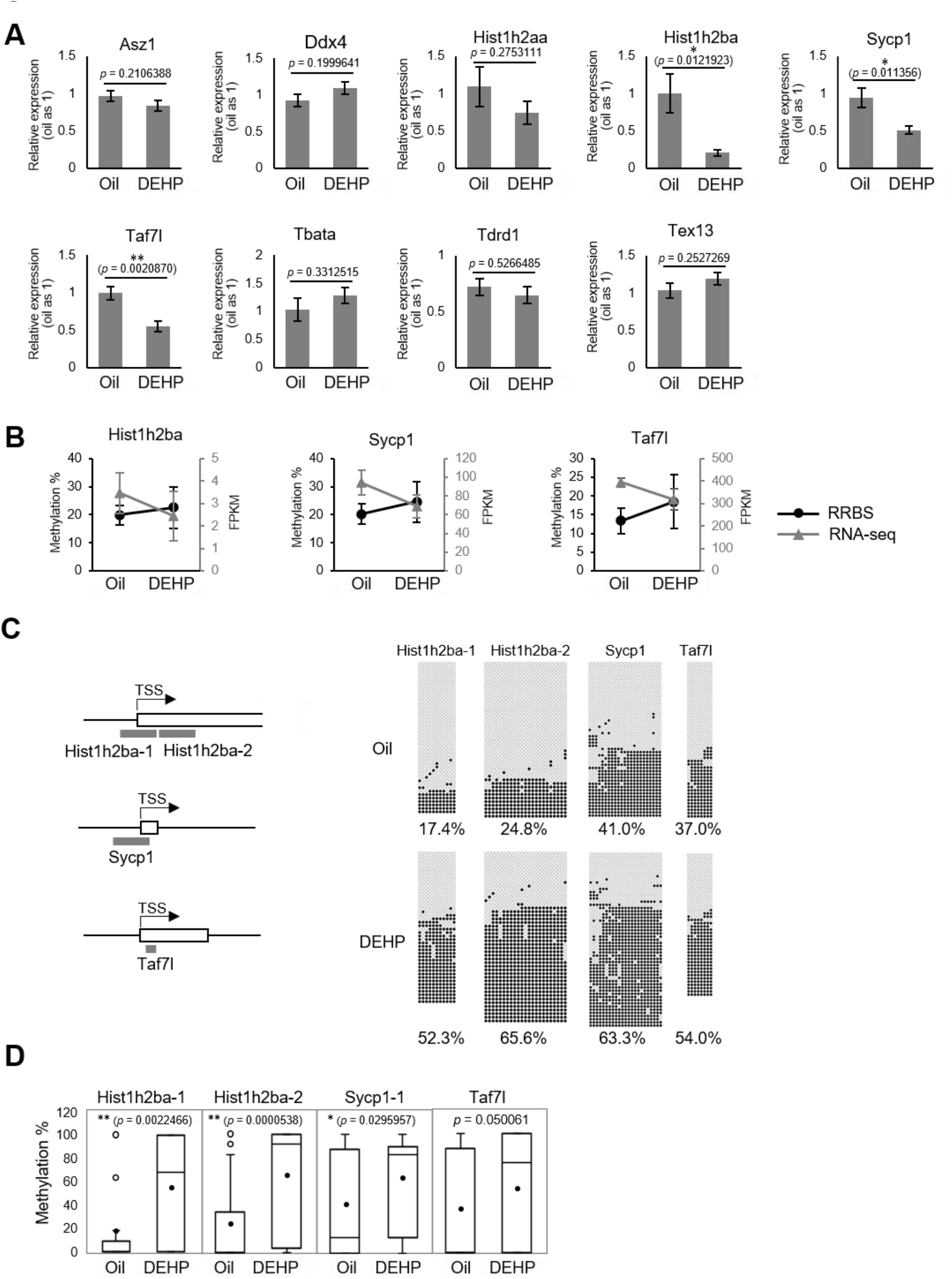
Gene expression and promoter methylation of spermatogenesis-related genes hypermethylated in both E19.5 germ cells and F1 spermatogonia following prenatal DEHP exposure. A. Relative mRNA levels in prenatally oil- or DEHP-treated adult F1 spermatogonia, as determined by RT-qPCR. Values of oil group are defined as 1.0. Values are plotted as mean ± SEM for samples obtained from six individuals. \**P* < 0.05, \*\**P* < 0.01 (unpaired two-sided Student’s t-test). B. Changes in promoter methylation (RRBS; black) and expression (RNA-seq; gray) of *Hist1h2ba, Sycp1*, and *Taf7l* following prenatal DEHP exposure in F1 spermatogonia. Error bars represent SEM (n=2). C. The regions detected in bisulfite sequencing of *Hist1h2ba, Sycp1*, and *Taf7l* are indicated in gray bars in the left panel. TSS represents the transcription start site. Boxes indicate the first exon. Methylation status of these regions in F1 spermatogonia obtained from four individuals are indicated in the right panel. Methylated and unmethylated CpGs are presented as closed circles and open circles, respectively. The percentage of methylated CpGs is indicated. D. Box-whisker plots of the CpG methylation levels shown in Fig 3C. The lines inside the boxes show the medians. The whiskers indicate the minimum and maximum values. Open and closed circles indicate outliers and mean value, respectively. Statistical analysis was performed using the Mann-Whitney U test.

We then focused on the 9 spermatogenesis-related genes that were hypermethylated in both E19.5 germ cells and F1 spermatogonia (Fig 2D); specifically, we used reverse transcription quantitative polymerase chain reaction (RT-qPCR) to examine the expression of these genes in F1 spermatogonia. This assay showed that the expression of *Hist1h2ba, Sycp1,* and *Taf7l* was significantly decreased in the DEHP group (compared to the oil group; Fig 3A). In agreement with this result, the RNA-seq data also showed downregulation of those genes in the DEHP-exposed group (compared to the oil group; Fig 3B). In addition, we confirmed that methylation levels of those genes identified by RRBS were increased by DEHP exposure (compared to those in the oil group; Fig 3B). We also examined the methylation levels of the promoter regions for *Hist1h2ba, Sycp1,* and *Taf7l* by bisulfite sequencing. As shown in Fig 3C and D, methylation of the CpG sites in two regions in the *Hist1h2ba* promoter and one region in *Sycp1* promoter were significantly elevated in DEHP-exposed spermatogonia (compared to the levels in the oil group). *Taf7l* in DEHP-exposed spermatogonia also tended to be hypermethylated, though this effect was not statistically significant. These results suggested that hypermethylation of *Hist1h2ba, Sycp1,* and *Taf7l* in F1 spermatogonia following maternal DEHP exposure contributes to the decreased expression of these genes, leading in turn to spermatogenic abnormalities. To explore the mechanism of hypermethylation, we examined expression levels of genes encoding DNA methyltransferases (DNMTs) using RNA-seq data. We observed that the *Dnmts* transcript levels (evaluated by their FPKM values) were altered by less than 1.5-fold, suggesting that any changes in methylation are not regulated by changes in *Dnmts* expression (figure supplement 4E).

Defects in spermatogenesis following maternal DEHP exposure also were observed in F2 testis. Therefore, we considered it likely that the hypermethylation of the spermatogenesis-related genes is maintained throughout spermatogenesis to transmit epigenetic traits to the F2 generation. However, none or few hypermethylated genes in E19.5 germ cells were shared between spermatocytes and round spermatids in F1 testis (Fig 2D), and methylation levels of the above-mentioned 3 spermatogenesis-related genes *(Hist1h2ba, Sycp1,* and *Taf7l)* in spermatogonia were decreased along with spermatogenesis (Fig 4). In addition, the methylation level in the DEHP group was decreased more drastically than was that in the oil group, achieving levels similar to those observed in round spermatids in the oil group. We also examined the expression and methylation of *Hist1h2ba, Sycp1,* and *Taf7l* in F2 spermatogonia. We found that F2 spermatogonia were not significantly altered in these parameters, but did exhibit a tendency to decreased expression of *Hist1h2ba, Sycp1,* and *Taf7l,* and to hypermethylation of *Hist1h2ba* and *Sycp1* in DEHP-treated groups (compared to control groups; figure supplement 5 A-C).

**Figure 4.**
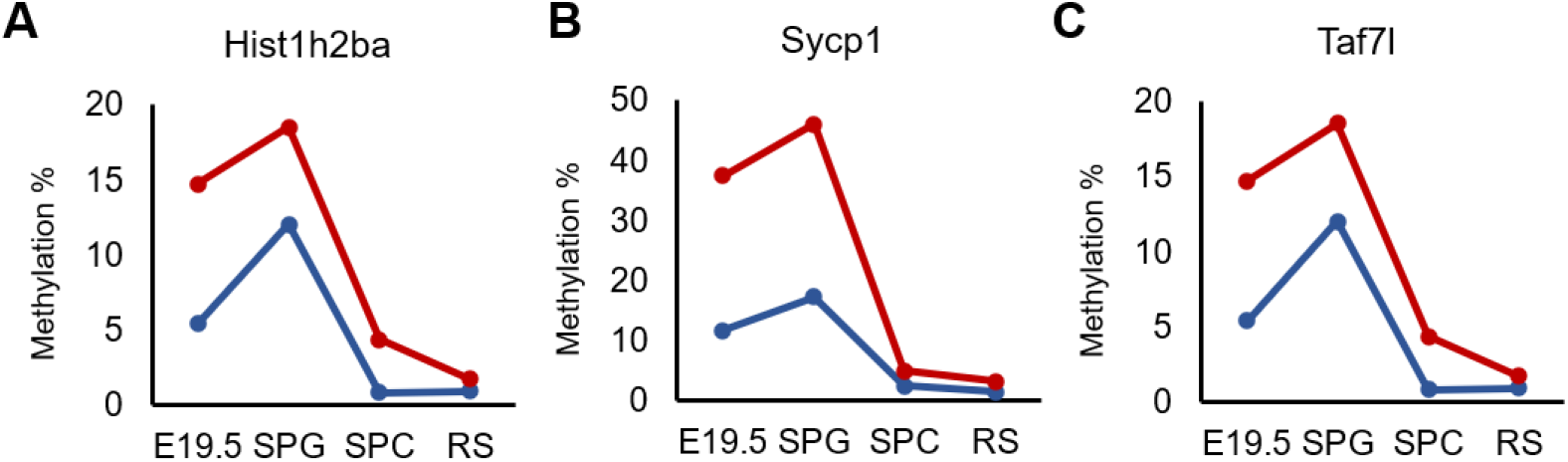
Changes in DNA methylation of the candidate epi-inutated genes in DEHP (red) and oil (blue) -treated E19.5 and adult F1 germ ceü populations. A-C. Promoter-methylation values (RRBS) *of Hist1h2ba* (A), *Sycp1* (B), and *Taf7l* (C) are plotted in each cell population.

We next examined the involvement of DNA methylation in regulation of the expression of *Hist1h2ba, Scyp1,* and *Taf7l.* Specifically, we employed a luciferase (Luc) reporter assay in cultured cells using promoter regions (−500 bp to +500 bp from the transcription start sites) of each of these genes (Fig 5A). We found that Luc activity was significantly decreased by methylation of the promoter of each gene (Fig 5B-D). The results supported the hypothesis that hypermethylation of *Hist1h2ba, Scyp1,* and *Taf7l* results in decreased expression in testicular germ cells.

**Figure 5.**
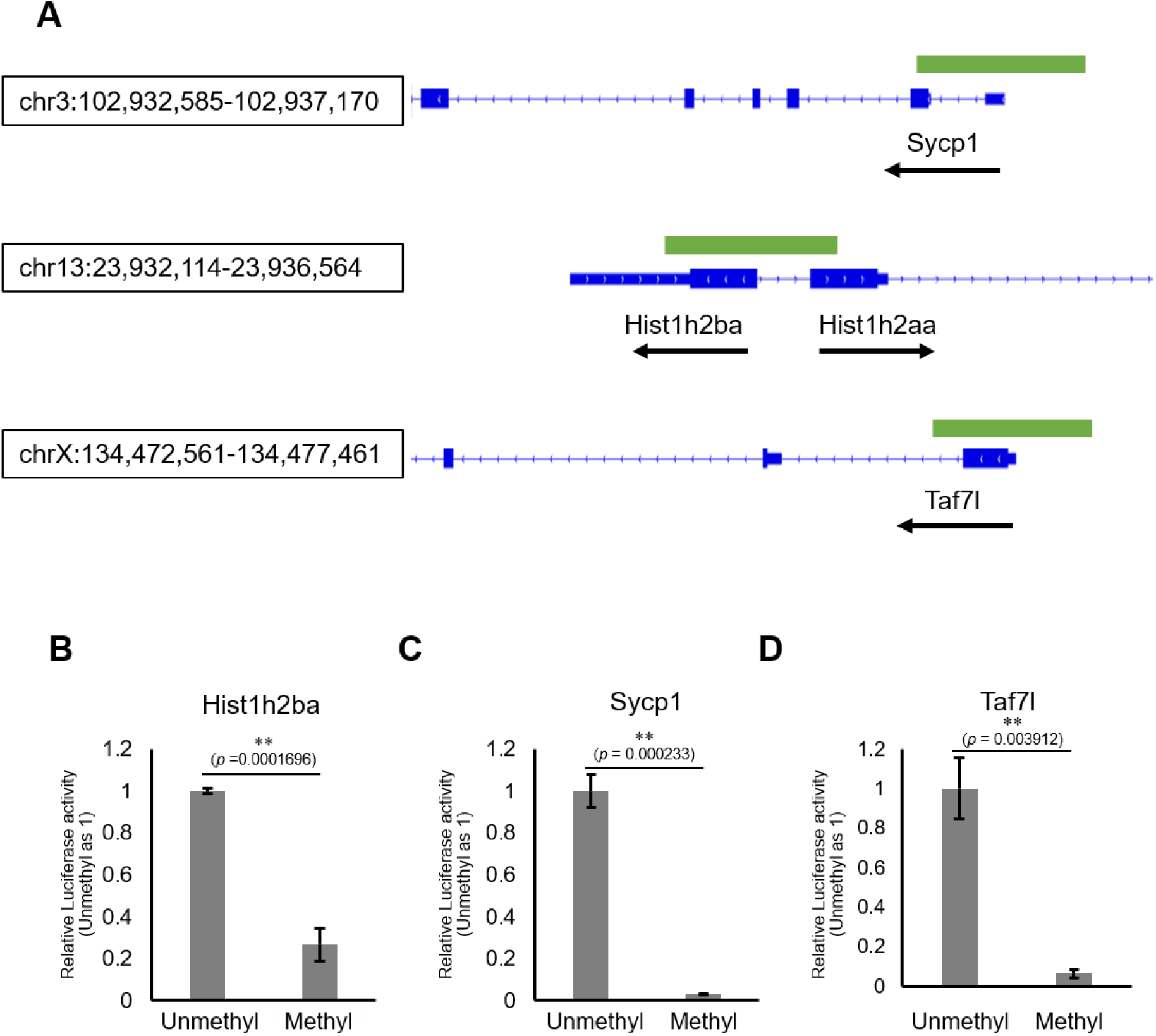
The effects on gene expression of promoter methylation in the candidate epi-mutated loci. A. Gene structures of *Hist1h2ba, Sycp1*, and *Taf7l*. Exons are indicated by blue bars, and the promoter regions (defined as ± 500 bp from TSS) used for firefly luciferase (Luc) assays are indicated by green bars. B-D. The Luc activity of the reporter vector with either methylated or unmethylated promoters of *Hist1h2ba* (B), *Sycp1* (C). and *Taf7l* (D) in HEK293T cells. Luc activity generated by the indicated vector was normalized to that generated by the *Renilla* phRL-TK vector. The Luc activity by the unmethylated vectors was defined as 1.0. Values are plotted as mean ± SEM of three independent experiments (n=3). \*\**P* < 0.01 (unpaired two-sided Student’s t-test).

## Discussion

### Close correlation between DNA methylation changes and abnormal spermatogenesis after maternal DEHP exposure

Induction of abnormal spermatogenesis in offspring following maternal DEHP exposure was first demonstrated using CD1 mice (Doyle *et al*., 2013), and subsequently was found in several strains of mice (including C57BL/6) and rats (Prados *et al*., 2015; Chen *et al*., 2015). In the present study, we also used C57BL/6 as well as Oct-4-deltaPE-GFP transgenic mice in a C57BL/6 genetic background to purify GFP-expressing fetal germ cells by flow cytometry. We first administered pregnant mice with 300 mg/kg DEHP, which is the same dose as used in a previous study with C57BL/6 (Prados *et al*., 2015); however, we obtained few pups under those conditions, presumably reflecting embryonic lethality at this dose of DEHP. We then tested different doses of DEHP, and identified dose levels at which we could obtain pups that showed germ cell abnormalities in F1 and F2 offspring and in F1 embryos that were similar to those reported previously (Doyle *et al*., 2013; Stenz *et al*., 2017; Ungewitter *et al*., 2017) at 100 or 150 mg/kg DEHP (Fig 1, figure supplement 1). Therefore, our experiments using C57BL/6 and Oct-4-deltaPE-GFP transgenic mice successfully reproduced the previously reported DEHP-induced histological abnormalities in testis, despite the use of a lower DEHP dose.

Using this experimental model, we identified a close correlation between DNA hypermethylation of spermatogenesis-related genes and their downregulation in F1 spermatogonia following maternal DEHP exposure (Fig 3). Previous reports have described promoter hypermethylation and downregulation of the *Svs3ab* gene, a gene that is involved in sperm motility, in the sperm of F1 males following maternal DEHP exposure (Stenz *et al*., 2017; Stenzid *et al*., 2019). Given that maternal DEHP exposure induces not only defective sperm but also abnormal spermatogenesis, we considered it likely that spermatogenesis-related genes are affected by DEHP (Fig 1, figure supplement 1); consistent with this hypothesis, we demonstrated promoter hypermethylation of three spermatogenesis-related gens, *Hist1h2ba, Sycp1,* and *Taf7l,* as well as downregulation of the expression of these three gene in the F1 spermatogonia following maternal DEHP exposure (Fig 2, Fig 3). In addition, these genes were hypermethylated in the E19.5 germ cells following maternal DEHP exposure (Fig 2), suggesting that DEHP is closely involved in the methylation of these loci. We also found that methylation of the promoters of these genes reduced their expression, as assessed by the methyl-luciferase assay (Fig 5), a result that implies the involvement of methylation of those genes in their downregulation. In a previous study, hypermethylation of some genes, but not that of spermatogenesis-related genes, was demonstrated in fetal male germ cells following maternal DEHP exposure (Prados *et al*., 2015); however, that study did not evaluate involvement of the hypermethylated genes in abnormal spermatogenesis. The inconsistency between our results and those of previous studies regarding hypermethylation of spermatogenesis-related genes by DEHP may be due, in part, to the use of different mouse strains. Specifically, our study employed mice with a C57BL/6 genetic background, as described above, while the previous study (which did not demonstrate testicular abnormalities) used mice with FVB and 129S1 genetic backgrounds (Iqbal *et al*., 2015). In addition, it has been reported that DEHP exposure does not cause any testicular abnormalities in FVB mice (Prados *et al*., 2015). Given that deficiencies in *Hist1h2ba, Sycp1,* or *Taf7l* result in abnormal spermatogenesis (as described below), our results imply that the downregulation of those genes following DEHP exposure is a cause of abnormal spermatogenesis.

### Effects of insufficient gene expression of Hist1h2ba, Scyp1, and Taf7l in spermatogenesis

The two histone variants HIST1H2aa and HIST1H2ba are abundant in the testis (Trostle-Weige *et al*., 1982), and male mice lacking both *Hist1h2aa* and *Hist1h2ba* are sterile (Shinagawa *et al*., 2015). Abnormal localization of nuclei in spermatids following maternal DEHP exposure (Fig 1A) is consistent with the phenotype observed in animals lacking both *Hist1h2ba* and *Hist1h2aa,* a phenotype that likely reflects impaired replacement of histones by protamine during spermiogenesis (Shinagawa *et al*., 2015). The *Hist1h2aa* and *Hist1h2ba* genes are located tandemly in inverted orientation on Chromosome 13 in mouse (Fig 5), and expression of the two loci is controlled by a shared promoter located between the two genes (Huh *et al*., 1991). Consistent with that structure, we found a tendency toward reduced *Hist1h2aa* expression in addition to significant downregulation of *Hist1h2ba* expression (Fig 3A) following maternal DEHP exposure, suggesting that hypermethylation of the shared promoter affects the expression of these two genes in testicular germ cells.

SYCP1 is a component of the synaptonemal complex, a structure that connects homologous chromosomes in meiotic prophase; *Sycp1* null mice are sterile because of spermatogenic arrest at the pachytene stage (de Vries *et al*., 2005). A similar abnormality was observed in F1 mice following maternal DEHP exposure (Fig 1A), a result that suggests that downregulation of *Sycp1* expression is a cause of abnormal spermatogenesis following maternal DEHP exposure. TAF7L is a testis-specific transcription-associated factor (Pointud *et al*., 2003). *Taf7l* null testis exhibits sperm with abnormal morphology and reduced motility, and large vacuoles are seen in the seminiferous tubules (Cheng *et al*., 2007); these defects resemble the abnormal tubules see in the F1 testis following maternal DEHP exposure. Taken together, these observations indicate that the *Hist1h2ba, Sycp1,* and *Taf7l* genes are epi-mutated following maternal DEHP exposure; the resulting downregulation of these gene may be a cause of abnormal spermatogenesis.

### Mechanism of hypermethylation of spermatogenesis-related genes in fetal germ cells following maternal DEHP exposure

Our study revealed that the spermatogenesis-related genes were hyper-methylated in fetal germ cells, while the methylation level of these genes was not significantly changed in testicular somatic cells following maternal DEHP exposure (Fig 2). These observations suggest a possible preference of DEHP-induced DNA methylation for germ cells and/or spermatogenesis-related genes. Previous studies have shown that maternal DEHP exposure induces DNA methylation not only in testis but also in other tissues such as adrenal gland and liver (Quinnies *et al*., 2015; Wen *et al*., 2020); notably, DEHP exposure resulted in a significant increase in the expression, in the liver of the offspring, of *Dnmt1,* a gene encoding a DNA methyltransferase. These results suggested that upregulation of *Dnmt1* by DEHP plays a role in DNA hypermethylation. However, we did not detect changes in *Dnmts* gene expression in fetal germ cells in the present work (figure supplement 4E). Therefore, it is likely that the induction (following DEHP exposure) of hypermethylation of spermatogenesis-related genes in germ cells may be mediated by other mechanisms.

Reactive oxygen species (ROS) may be involved in DEHP-induced DNA methylation, given that mono (2-ethylhexyl) phthalate (MEHP), a hydrolyzed metabolite of DEHP, induces ROS production in adult testicular germ cells (Kasahara *et al*., 2002), and that ROS directly enhances DNA methylation (Afanas’ev, 2014). In fetal germ cells, the chromatin structure around spermatogenesis-related genes is highly accessible (Li *et al*., 2018), and may facilitate preferential DNA methylation of spermatogenesis-related genes in fetal germ cells in response to the accumulation of ROS following DEHP exposure. Alternatively, DEHP-or MEHP-binding molecules may be involved in the observed DNA methylation. DEHP and MEHP are ligands of peroxisome proliferator-activated receptors (PPARs) and the constitutive androstane receptor (CAR); these receptors regulate downstream pathways (Corton & Lapinskas, 2005; Eveillard *et al*., 2009), and may thereby promote DNA methylation via unknown mechanisms, including recruitment of DNA methylation-related molecules to target genes. Those possibilities await further investigation.

### Possible mechanisms of intergenerational inheritance of abnormal spermatogenesis following maternal DEHP exposure

The expression and methylation of *Hist1h2ba, Sycp1,* and *Taf7l* in F2 spermatogonia tended to be slightly affected by maternal DEHP exposure (figure supplement 5). In our results, methylation levels in these genes in F2 spermatogonia generally were lower than those in F1 spermatogonia, even in the oil-treated group (Fig 3, figure supplement 5B, C); these differences may be due, at least in part, to differences in the purity of the spermatogonia. We fund that enrichment of *Gfra1* expression in sorted spermatogonia fractions were higher in those of F1 than those of F2, though its enrichment between oil- and DEHP-treated groups of F1 and of F2 were comparable (figure supplement 5D). Because the F2 alleles are inherited from DEHP-exposed males and untreated females, the influence of DEHP-induced epigenetic changes reasonably may become weaker in the F2 alleles compared with those in the F1 alleles. Additionally, those genes became hypomethylated during spermatogenesis in F1 testis (Fig 4), suggesting that the hypermethylation of the genes by DEHP, if any, is not directly transmitted to the F2. A recent study suggested that binding of transcription factors (TFs) at CpG sites affects the methylation status of these sites during demethylation in PGCs and affect subsequent re-methylation of these sites; notably, CpG sites not bound by TFs could be remethylated after demethylation (Kremsky & Coreces, 2020). These results imply that *Hist1h2ba, Sycp1,* and *Taf7l* are demethylated during spermatogenesis, and subsequently are re-methylated at some developmental time point(s) after fertilization.

In the case of embryonic undernutrition, abnormal DNA hypomethylation in F1 sperm is not inherited in F2 sperm and somatic tissues; however, the transcription levels of several genes near the regions with altered DNA methylation in F1 sperm are dysregulated in F2 somatic tissues (Radford *et al*., 2014). These results suggest that changes in DNA methylation perturb (indirectly) gene expression patterns in subsequent generations. Modified histones that are bound to specific genomic regions, such as H3K4 trimethylation (me3) bound at CpG-rich promoters and H3K9me3 bound at satellite repeats in the sperm genome, are not replaced by protamine during sperm development (Yamaguchi *et al*., 2018). Maternal undernutrition, as well as maternal DEHP exposure, may affect such histone modifications in F1 sperm in a DNA methylation status-dependent manner, which in turn may influence phenotypes in the F2. Further work will be required to explore this possibility.

## Materials and Methods

### Animals and DEHP treatment

C57BL/6J mice were purchased from Japan SLC. Oct4-deltaPE-GFP transgenic mice (Yoshimizu *et al*., 1999) were maintained in a C57BL/6J genetic background. The mice were maintained and bred in an environmentally controlled and specific-pathogen-free facility, the Animal Unit of the Institute of Development, Aging and Cancer (Tohoku University), according to the guidelines for experimental animals defined by the facility. Animal protocols were reviewed and approved by the Tohoku University Animal Studies Committee. Maternal DEHP exposure was performed as described previously (Stenz *et al*., 2017). Briefly, female C57BL/6J mice (8 to 12 weeks old) were mated with male C57BL/6J or Oct4-deltaPE-GFP transgenic mice; noon on the day of mucus plug observation was defined as E0.5. Prenatal exposure to DEHP was performed by oral (gavage) administration of pregnant female mice, performed once per day from E8 to E18. At each administration, 200 μL of either corn oil (Sigma C8267) (for controls) or 200 μL of DEHP (Sigma 80030) diluted in corn oil to provide a dose of 100 mg/kg/day or 150 mg/kg/day for C57BL/6J or Oct4-deltaPE-GFP males, respectively, was delivered by oral gavage. The treated gestating dams were considered the F0 generation (F0), and the pups born from the F0 dams were considered the F1 generation (F1) offspring. The F2 generation was obtained by breeding males of the F1 of DEHP-treated animals or vehicle control animals with nontreated C57BL/6J females. We analyzed three to six mice of each conditions as biological replicates to statistically evaluate the effect of DEHP.

### Purification of germ cells

E19.5 embryos were obtained from DEHP- or corn oil-treated female mice; embryonic testes were isolated, and the albuginea was removed in Dulbecco’s modified Eagle medium (DMEM, Gibco 11965-092) containing 10% fetal bovine serum (FBS) (Biosera FB-1380). Testes then were incubated for 15 min at 37 °C in phosphate-buffered saline (PBS) containing 0.5 mg/mL trypsin and 0.2 mg/mL EDTA (Sigma T4174). The cells were dissociated by pipetting, and then were filtered through a 40-μm-pore Cell strainer (BD Falcon 352340). A Bio-Rad S3e cell sorter was used to purify viable germ cells with intense Oct4-deltaPE-GFP expression and testicular somatic cells (Somas) without Oct4-deltaPE-GFP expression. For DNA isolation, one-quarter-volume of sorted cell suspension was centrifuged and stored at −80 °C after removal of the supernatant. For RNA isolation, the remainder of the cell suspension was centrifuged, resuspended in buffer RLT (Qiagen 79216) containing 1% beta-mercaptoethanol, and stored at −80 °C. For each cell sorting, we obtained 1 to 16 embryos. The cells obtained from 4 independent sorting were combined before DNA or RNA extraction, which yielded suspensions containing approximately 12000 cells for DNA isolation and 40000 cells for RNA isolation. We repeated four more sorting and combined the sorted cells to obtain a biological replicate. Dissociation of testicular cells from adult mice and subsequent staining were carried out as described previously (Bastos *et al*., 2005; Ota *et al*., 2019). Testes were dissected from F1 or F2 mice at approximately postnatal day (P) 200. After the albuginea was removed, testes were incubated at 32 °C for 25 min in 6 mL of Gey’s Balanced Salt Solution (GBSS; Sigma-Aldrich G9779) containing 1.2 mg/mL of Collagenase Type I (Sigma-Aldrich C0130), and the seminiferous tubules were dissociated. Interstitial cells were removed by filtration with the Cell strainer. Seminiferous tubules retained on the filter were collected and incubated at 32 °C for 25 min in GBSS containing 1.2 mg/mL of Collagenase Type I and 5 μg/mL DNase (Roche 11284932). Cell aggregates were sheared gently by 10 rounds of pipetting with a wide orifice plastic transfer pipet and filtered through the Cell strainer to remove cell clumps. Cells were washed with GBSS and then resuspended in GBSS containing 1% FBS. Twenty million cells were diluted in 2 mL of GBSS containing 1% FBS and stained with 5 μg/mL of Hoechst 33342 (Invitrogen H3570) for 1 h at 32 °C. Cells were kept on ice and protected from light until sorting. Before sorting, 0.25 μg/mL of propidium iodide (BD 51-66211E) was added to the stained cells, and the mixture was filtered through the Cell strainer. The cells were sorted using a Becton-Dickinson FACSAria II cell sorter, and spermatogonia, spermatocytes, and round spermatids were collected according to their staining patterns. Confirmation of the purity of the sorted germ cells was evaluated by the assessing the expression of stage-specific germ cell marker genes as follows: *Gfra1* for spermatogonia (Hoffman *et al*., 2005), *Scp3* for spermatocytes (Lammers *et al*., 1994), and *Acrv1* for spermatids (Reddi *et al*., 1995). We obtained two biological replicates for each cell type in F1 testes for transcriptome and methylome; one was composed of the cells from one individual, and the other was composed of pooled cells from two individuals to secure enough number of cells. Each sample was further divided for transcriptome (approximately 50000 cells) and methylome (approximately 15000 cells) analyses. For RT-qPCR and bisulfite sequencing, spermatogonia collected from one individual were used as one biological replicate.

### Histological examination

Testes from E19.5 embryos of the F1 generation and P200 male mice of the F1 and F2 generations were fixed overnight at 4 °C in <u>Bouin’s</u> solution with rotation, and then were embedded in paraffin. Five-micrometer-thick serial sections were cut and mounted on 3-aminopropyltriethoxysilane (APS) -coated slides (Matsunami APS-01); the mounted sections were deparaffinized, stained with hematoxylin and eosin, and examined using an optical microscope (Leica). Digital images were obtained using a LAS4.4 (Leica), and abnormal tubules were determined by the histological criteria described in the Results section. To determine the percentage of abnormal tubules, the numbers of abnormal tubules and total tubules were counted in each testicular section. Four sections, each 100 μm apart in a given testis, were counted.

### Reduced representation bisulfite sequencing (RRBS)

The purified cells described above were suspended in a lysis buffer (0.14 mM β-mercaptoethanol, 0.24 mg/ml Proteinase K, 150 mM NaCl, 10 mM Tris-HCl [pH 8.0], 10 mM EDTA [pH 8.0], and 0.1% SDS) and incubated at 55 °C for 2 hours. Genomic DNA then was isolated using phenol/chloroform extraction and ethanol precipitation. RRBS libraries were generated as previously reported (Gu *et al*., 2011). Briefly, 20 ng of genomic DNA was subjected to MspI digestion (NEB, Beverly, MA, USA), with subsequent end repair/dA-tailing reaction using Klenow Fragment (3’-5’ exo-) (NEB M0212S) and ligation with Illumina sequencing adapters using T4 DNA ligase (NEB). The resulting mixture was electrophoresed in NuSieve 3:1 agarose (Lonza 50091), and fragments sized 150-350 bp were excised and then purified using a MinElute Gel Extraction kit (QIAGEN 28604). The purified fragments were treated with sodium bisulfite using an EZ DNA Methylation-Gold Kit (Zymo Research D5005). Library amplification and indexing were performed with KAPA HiFi HotStart Uracil+ ReadyMix (2×) (Kapa Biosystems KK2801). The PCR amplification was carried out as follows: an initial denaturation at 95 °C for 2 min; 13 cycles at 98 °C for 20 sec, 65 °C for 30 sec, and 72 °C for 30 sec; and a final 1-min extension at 72 °C. The RRBS libraries were purified using Agencourt AMPure XP (Beckman Coulter A63880), quantified with a Kapa Library Quantification Kit (Kapa Biosystems KK4824), and sequenced on an HiSeq 2500 platform (Illumina) with 100-bp single-end reads using a TruSeq SR Cluster Kit v3-cBot-HS and TruSeq SBS Kit v3-HS (Illumina). Sequenced reads were processed using an Illumina standard base-calling pipeline (v1.8.2), and the index and adapter sequences were removed. The first and last 4 bases were trimmed, and the resulting reads were aligned to Mouse Genome Build 37 (mm10) using Bismark (v.0.10.1) (Krueger & Andrews, 2011) with default parameters. The methylation level of each cytosine was calculated using the Bismark methylation extractor. We analyzed only CpG cytosines covered with 3 reads. Annotations of Refseq genes and repeat sequences were downloaded from the UCSC Genome Browser. Refseq genes encoding microRNAs and small nucleolar RNAs as well as mitochondrial DNA were excluded from our analyses. Promoters were defined as regions 500 bp upstream and downstream from transcription start sites (TSSs) of Refseq transcripts. Gene bodies were defined as transcribed regions of Refseq transcripts except for promoters. For calculation of the mean methylation levels, we analyzed promoters containing ≧10 CpG cytosines with sufficient coverage for calculation of the methylation levels. We analyzed two biological replicates to show reproducibility.

### RNA sequencing

RNA-seq libraries were prepared from total RNA purified from sorted germ cells with a RNeasy Micro Kit (QIAGEN 74004). The libraries were clonally amplified on a flow cell and sequenced on HiSeq2500 (HiSeq Control Software v2.2.58, Illumina) with 51-mer single-end sequences. Image analysis and base calling were performed using Real-Time Analysis Software (v1.18.64, Illumina). For gene expression analysis, reads were mapped to the mouse genome (UCSC mm10 genome assembly and NCBI RefSeq database) using TopHat2 and Bowtie. Cufflinks was used to estimate gene expression levels based on fragments per kilobase of exon per million mapped reads (FPKM) normalization. Differentially expressed genes (DEGs) were extracted from the Cuffdiff results. We analyzed two biological replicates to show reproducibility, except E19.5 DEHP sample due to failed library preparation.

### Data analysis

The Database for Annotation, Visualization, and Integrated Discovery (DAVID, https://david.ncifcrf.gov/, Classification stringency: medium) (Huang *et al*., 2009) was used for functional annotation. Venn diagram analysis was performed by InteractiVenn (http://www.interactivenn.net/). Heatmaps were created with MetaboAnalyst 5.0 (http://www.metaboanalyst.ca). Chromosome positions of hyper-or hypo-methylated genes were visualized by Ensembl (https://asia.ensembl.org/index.html).

### Bisulfite sequencing analysis

Genomic DNA was isolated using phenol/chloroform extraction and ethanol precipitation. Bisulfite conversion of the DNA was performed using EZ DNA Methylation-Gold Kit (ZYMO Research D5005). Nested PCR was performed using EpiTaq HS DNA Polymerase (TaKaRa R110A). The sequences of the PCR primer were designed with MethPrimer (http://www.urogene.org/cgi-bin/methprimer/methprimer.cgi) (Li & Dahiya, 2002). The primer sequences are shown in Supplementary Table 5. The PCR products were gel-purified, sub-cloned into the pGEM-T Easy vector (Promega A1360), and sequenced using an ABI PRISM 3100-Avant Genetic Analyzer (Applied Biosystems). Sequence data were analyzed with Quantification tool for Methylation Analysis (http://quma.cdb.riken.jp/top/quma_main_j.html) (Kumaki *et al*., 2008). The data were obtained from four and three animals in each treatment group of F1 and F2, respectively, to evaluate statistically significant differences and data for 6-18 clones in each animal were used for analysis.

### Quantitative RT-PCR

Total RNA was extracted from cells using a RNeasy Micro Kit according to the manufacturer’s instructions. RNAs were reverse-transcribed using SuperScript III reverse transcriptase (Invitrogen 18080093) and random primers (Promega C1181). Real-time PCR was performed using Power SYBR Green PCR Master Mix (Applied Biosystems 4367659). Thermal conditions were 2 min at 50 °C, 10 min at 95 °C, and 45 cycles of 15 sec at 95 °C and 60 sec at 60 °C. Sequences of the primers used for the PCR reaction are shown in Supplementary Table 5. The *Arbp* transcript was used as an internal control. The relative expression was analyzed using comparative CT method. The data were obtained from six and three animals in each treatment group of F1 and F2, respectively, to evaluate statistically significant differences.

### Luciferase reporter assay

The gene regions between 500 bp upstream and downstream from the TSSs of *Hist1h2ba, Sycp1,* and *Taf7l* were amplified and sequenced. Each sequence was cloned into the CpG-free pCpGL-basic Luciferase vector (Klug & Rehli, 2006). Luciferase reporter constructs were either mock-treated or methylated in vitro with SssI CpG methyltransferase for 4 h at 37 °C and purified with the QIAquick Purification Kit (QIAGEN 28704). Reporter plasmid (500 ng) and *Renilla* phRL-TK control vector (50 ng; Promega E2241) were cotransfected into HEK293T cells cultured in DMEM containing 10% FBS using Lipofectamine LTX Reagent with PLUS Reagent (Invitrogen 15338100). After 48 h, cells were lysed, and the relative luciferase activities were analyzed using the Dual-Luciferase Reporter Assay System (Promega E1910) on a Lumat LB 9507 (Berthold). Firefly luciferase (Luc) activity of individual transfections was normalized against *Renilla* luciferase activity. We analyzed data from three independent experiments to evaluate statistically significant differences.

### Statistical analysis

The significance of difference was assessed by the unpaired two-sided Student’s t test except for bisulfite sequence. Statistical analysis of DNA methylation levels was performed by using the Mann- Whitney U test. The level of significance was set at P-value < 0.05.

## Data Availability

RNA-seq data: pending

RRBS data: pending

## Acknowledgements

The authors thank Dr. D. Okamura and K. Mochizuki for helpful advices and discussions, all the members of Cell Resource Center for Biomedical Research for helpful discussions, the Biomedical Research Core of Tohoku University Graduate School of Medicine and the Center of Research Instruments of Institute of Development, Aging and Cancer (IDAC), Tohoku University for use of instruments and technical supports. YT was supported by a Grant-in-Aid for Japan Society for the Promotion of Science (JSPS) Fellows (18J40019). YM. was supported by Grant-in-Aid for Scientific Research (KAKENHI) (B) (grant 19H03231) from the Ministry of Education, Culture, Sports, Science and Technology of Japan (MEXT), and AMED-CREST (grant #JP17gm0510017h) from the Japan Agency for Medical Research and Development.

## Author contributions

Conceptualization: YT, YM; Data curation: YT, YM; Funding acquisition: YT, YM; Investigation: YT, HH; Methodology: YT, HH; Project administration: YM, TA; Resources: YT, AT, YI-M; Supervision: YM; Visualization: YT, YM; Writing: YT, YM

---

Supplementary File 1. Methylation levels (%) of various genomic regions.

Supplementary File 2. List of hyper- (DEHP-oil (%) > 5) and hypo- (DEHP-oil (%) < −5) methylated genes in E19.5 testicular germ cells and somatic cells.

Supplementary File 3. List of hyper- (DEHP-oil (%) > 5) and hypo- (DEHP-oil (%) < −5) methylated genes in spermatogonia, spermatocyte, and round spermatid.

Supplementary File 4. List of down- and up-regulated genes in E19.5 testicular germ cells, and spermatogonia, spermatocyte, and round spermatid in adult testis (more than 2-Fold-Changes).

Supplementary File 5. Primers used in this study.

**figure supplement 1.**
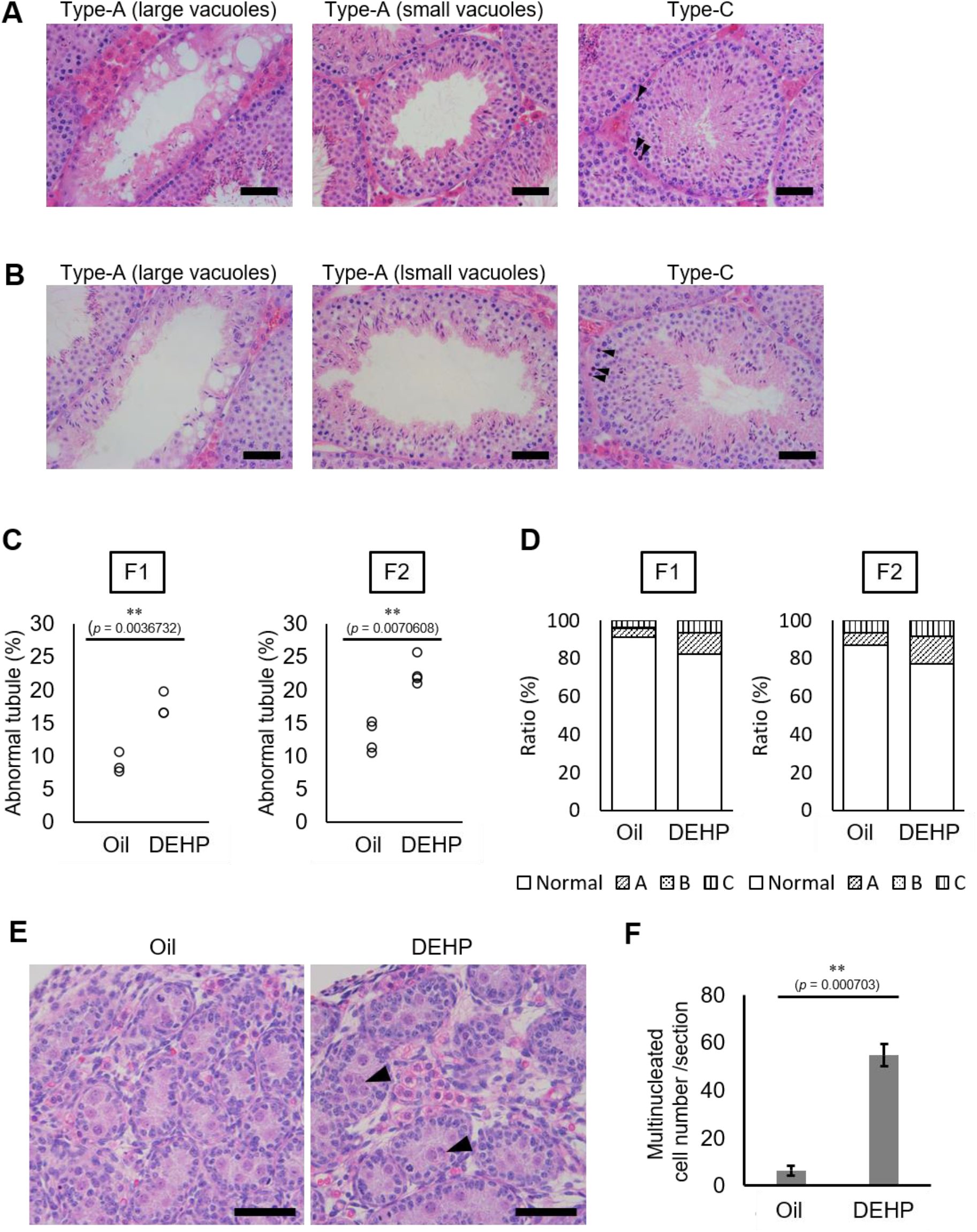
Histological analysis of the testicular tubules of prenatally DEHP-exposed offspring from C57BL/6 females mated with Oct4-deltaPE-GFP transgenic males. A, B. Representative images of testicular tubules in F1 (A) and F2 (B). Abnormal testicular tubules were categorized as “Ttype A” (presence of various size of vacuoles) and ‘‘Type C” (presence of dead cells; indicated by arrowheads). “Type-B” (no lumen and complete loss of germ cell organization) abnormal tubules were very rare in F1 and absent in F2. Arrowheads indicate apoptotic cells. C. Ratios of abnormal tubules in F1 and F2 testes. (F1: n=3; F2: n=4). D. Ratios of abnormal tubule types in F1 and F2. E. Representative histology of testis from oil- or DEHP-treated E19.5 embryos. Multinucleated cells are indicated by arrowheads. F. Quantification of the multinucleated cell number per section. Values are plotted as mean ± SEM of all serial sections from one testis in each of three independent embryos. ** *P* < 0.01 (unpaired two-sided Student’s t-test). Scale bars: 50 μm.

**figure supplement 2.**
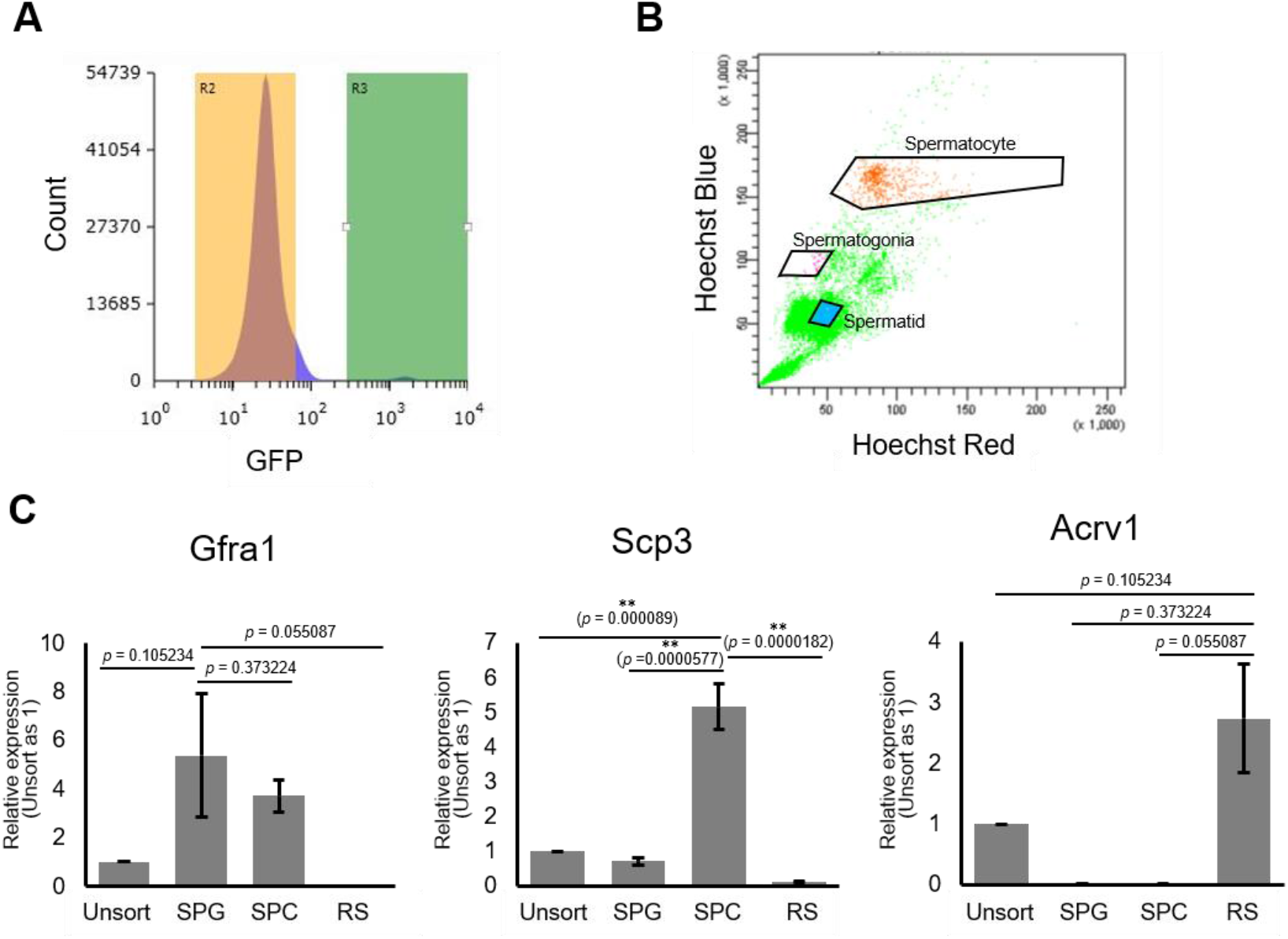
Purification of germ cells from E19.5 and adult testis. A. A representative FACS histogram of E19.5 testis. GFP-positive germ cells in the R3 gate (green color), and GFP-negative testicular somatic cells in the R2 gate (yellow color), were collected. B. A representative FACS plot of F1 testicular cells stained with Hoechst 33342. Gates for spermatogonia, spermatocytes, and round spermatids are indicated. C. D. Relative expression of stage-specific germ cell marker genes in F1 (C) sorted cells was determined by RT-qPCR., and consisted of the following: *Gfra1* for spermatogonia, *Scp3* for spermatocytes, and *Acrv1* for spermatids. SPG; spermatogonia, SPC; spermatocytes. RS; round spermatids. Values were plotted as mean ± SEM of each cell sample from 6 individuals of F1. \*\**P* < 0.01 (unpaired two-sided Student’s t-test).

**figure supplement 3.**
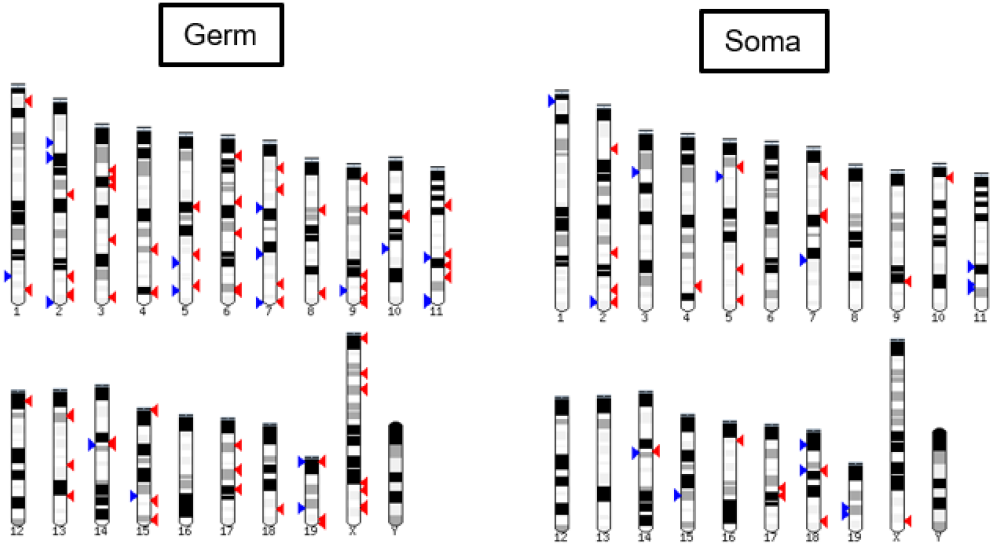
Methylome of prenatally DEHP-exposed testicular cells. Chromosomal positions of differentially promoter-methylated genes in E19.5 germ and somatic cells. Red arrowhead: more than 5% increase in methylation in DEHP group compared to oil group; blue arrowhead: more than 5% decrease in methylation in DEHP group compared to oil group.

**figure supplement 4.**
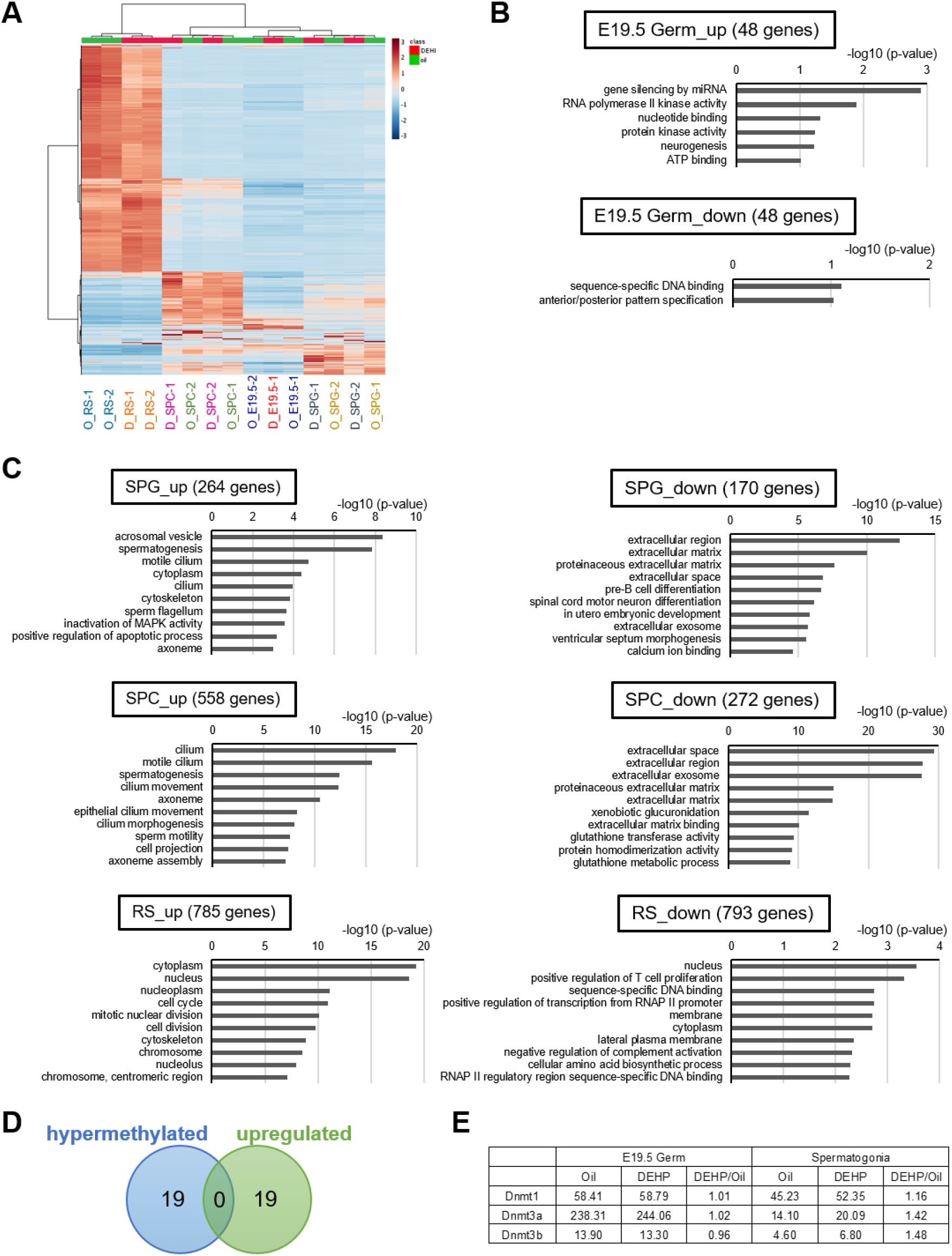
Transcriptome of testicular germ cells after prenatal DEHP exposure. A. A heatmap of FPKM values in E19.5 and adult germ cell populations of F1 prenatally exposed to oil (O) or DEHP (D). B, C. Functional annotations of genes exhibiting up- or down-regulation (more than 2-fold changes) following DEHP exposure (compared to oil exposure) in E19.5 germ cells (B) and F1 germ cell populations. SPG: spermatogonia; SPC: spermatocytes; RS: round spermatids. D. Venn diagram analysis of genes in spermatogenesis GO in upregulated or hypermethylated by DEHP in spermatogonia. E. FPKM values and fold changes in the expression of *Dnmtl, Dnmt3a,* and *Dnmt3b* in E19.5 germ cells and adult F1 spermatogonia.

**figure supplement 5.**
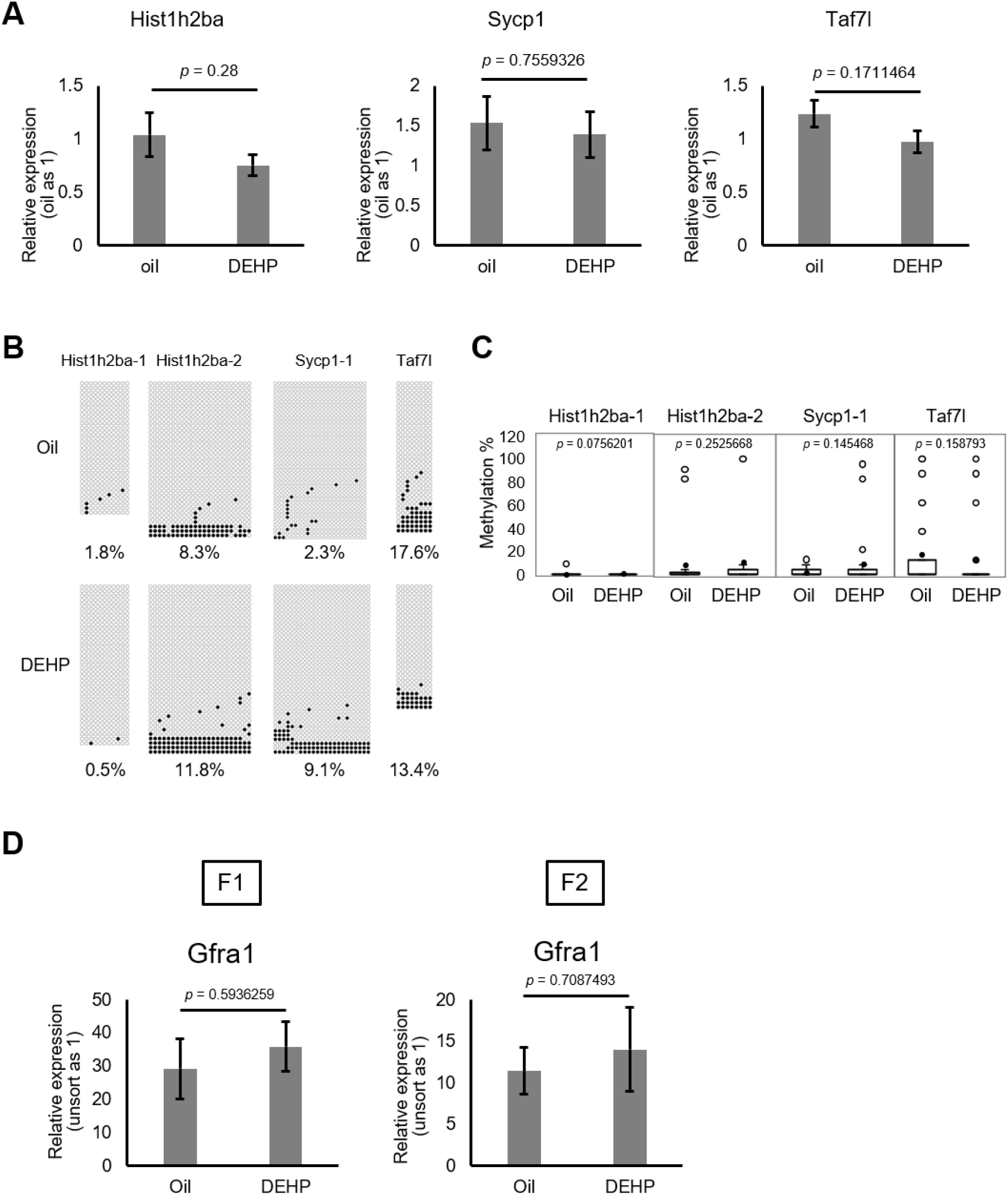
Gene expression and promoter methylation of *Hist1h2ba, Sycp1,* and *Taf7l* in F2 spermatogonia. A. Relative mRNA levels determined by RT-qPCR in prenatal oil- or DEHP-treated adult F2 spermatogonia. Values are plotted as mean ± SEM of spermatogonia samples obtained from three individuals. B. Methylation status of these regions determined by bisulfite sequencing in spermatogonia obtained from three individuals. Methylated and unmethylated CpGs are presented as closed circles and open circles, respectively. The percentage of methylated CpGs is indicated. C. Box-whisker plots of the CpG methylation levels shown in figure supplement 5B. The lines inside the boxes show the medians. The whiskers indicate the minimum and maximum values. Open and closed circles indicate outliers and mean value, respectively. Statistical analysis was performed using the Mann-Whitney U test. D. Relative expression of *Gfra1* in F1 and F2 sorted spermatogonia determined by RT-qPCR. Values were plotted as mean ± SEM of each cell sample from 4 individuals of F1 and 3 individuals of F2. Statistics utilized in A and D were unpaired two-sided Student’s t-test.

## References

Afanas’ev I (2014) New nucleophilic mechanisms of ROS-dependent epigenetic modifications: Comparison of aging and cancer. Aging Dis 5: 52–62

Anway MD, Cupp AS, Uzumcu M & Skinner MK (2005) Epigenetic Transgenerational Action of Endocrine Disruptors and Male Fertility. Science 308: 1466–9

Bastos H, Lassalle B, Chicheportiche A, Riou L, Testart J, Allemand I & Fouchet P (2005) Flow cytometric characterization of viable meiotic and postmeiotic cells by Hoechst 33342 in mouse spermatogenesis. Cytom Part A 65A: 40–49

Brehm E, Rattan S, Gao L & Flaws JA (2018) Prenatal exposure to Di(2-ethylhexyl) phthalate causes longterm transgenerational effects on female reproduction in mice. Endocrinology 159: 795–809

Carone BR, Fauquier L, Habib N, Shea JM, Hart CE, Li R, Bock C, Li C, Gu H, Zamore PD, et al (2010) Paternally induced transgenerational environmental reprogramming of metabolic gene expression in mammals. Cell 143: 1084–1096

Chen J, Wu S, Wen S, Shen L, Peng J, Yan C, Cao X, Zhou Y, Long C, Lin T, et al (2015) The mechanism of environmental endocrine disruptors (DEHP) induces epigenetic transgenerational inheritance of cryptorchidism. PLoS ONE 10: e0126403

Cheng Y, Buffone MG, Kouadio M, Goodheart M, Page DC, Gerton GL, Davidson I & Wang PJ (2007) Abnormal Sperm in Mice Lacking the Taf7l Gene. Mol Cell Biol 27: 2582–2589

Corton JC & Lapinskas PJ (2005) Peroxisome proliferator-activated receptors: Mediators of phthalate ester-induced effects in the male reproductive tract? Toxicol Sci 83:4–17

Daxinger L & Whitelaw E (2012) Understanding transgenerational epigenetic inheritance via the gametes in mammals. Nat Rev Genet 13: 153–162

Doyle TJ, Bowman JL, Windell VL, McLean DJ & Kim KH (2013) Transgenerational effects of Di-(2-ethylhexyl) phthalate on testicular germ cell associations and spermatogonial stem cells in mice. BiolReprod 88: 112

Eveillard A, Mselli-Lakhal L, Mogha A, Lasserre F, Polizzi A, Pascussi JM, Guillou H, Martin PGP & Pineau T (2009) Di-(2-ethylhexyl)-phthalate (DEHP) activates the constitutive androstane receptor (CAR): A novel signalling pathway sensitive to phthalates. Biocheml Pharmacol 77: 1735–1746

Gu H, Smith ZD, Bock C, Boyle P, Gnirke A & Meissner A (2011) Preparation of reduced representation bisulfite sequencing libraries for genome-scale DNA methylation profiling. Nature Protocols 6: 468–481

Hofmann MC, Braydich-Stolle L & Dym M (2005) Isolation of male germ-line stem cells; influence of GDNF. Developmental Biology 279: 114–24

Huang DW, Sherman BT & Lempicki RA (2009) Systematic and integrative analysis of large gene lists using DAVID bioinformatics resources. Nate Protoc 4: 44–57

Huh NE, Hwang IW, Lim K, You KH & Chae CB (1991) Presence of a bi-directional S phase-specific transcription regulatory element in the promoter shared by testis-specific TH2A and TH2B histone genes. Nucleic Acids Res 19: 93–98

Iqbal K, Tran DA, Li AX, Warden C, Bai AY, Singh P, Wu X, Pfeifer GP & Szabó PE (2015) Deleterious effects of endocrine disruptors are corrected in the mammalian germline by epigenome reprogramming. Genome Biol 16: 59

Kasahara E, Sato EF, Miyoshi M, Konaka R, Hiramoto K, Sasaki J, Tokuda M, Nakano Y & Inoue M (2002) Role of oxidative stress in germ cell apoptosis induced by di(2-ethylhexyl) phthalate. Biochem J 365(Pt 3): 849–56

Klug M & Rehli M (2006) Functional analysis of promoter CpG methylation using a CpG-free luciferase reporter vector. Epigenetics 1: 127–130

Kremsky I & Corces VG (2020) Protection from DNA re-methylation by transcription factors in primordial germ cells and pre-implantation embryos can explain trans-generational epigenetic inheritance. Genome Biol 21: 118

Krueger F & Andrews SR (2011) Bismark: A flexible aligner and methylation caller for Bisulfite-Seq applications. Bioinformatics 27: 1571–1572

Kumaki Y, Oda M & Okano M (2008) QUMA: quantification tool for methylation analysis. Nucleic Acids Res 36(Web Server issue): W170–175

Lammers JH, Offenberg HH, van Aalderen M, Vink AC, Dietrich AJ & Heyting C (1994) The gene encoding a major component of the lateral elements of synaptonemal complexes of the rat is related to X-linked lymphocyte-regulated genes. Mol Cell Biol 14: 1137–1146

Li J, Shen S, Chen J, Liu W, Li X, Zhu Q, Wang B, Chen X, Wu L, Wang M, et al (2018) Accurate annotation of accessible chromatin in mouse and human primordial germ cells. Cell Res 28: 1077–1089

Li LC & Dahiya R (2002) MethPrimer: designing primers for methylation PCRs. Bioinformatics 18(11):1427–31

ben Maamar M, Nilsson E, Sadler-Riggleman I, Beck D, McCarrey JR & Skinner MK (2019) Developmental origins of transgenerational sperm DNA methylation epimutations following ancestral DDT exposure. Dev Biol 445: 280–293

Ng SF, Lin RCY, Laybutt DR, Barres R, Owens JA & Morris MJ (2010) Chronic high-fat diet in fathers programs β 2-cell dysfunction in female rat offspring. Nature 467: 963–966

Ota H, Ito-Matsuoka Y & Matsui Y (2019) Identification of the X-linked germ cell specific miRNAs (XmiRs) and their functions. PLoS ONE 14: e0211739

Trostle-Weige PK, Meistrich ML, Brock WA, Nishioka K & Bremer JW (1982) Isolation and characterization of TH2A, a germ cell-specific variant of histone 2A in rat testis. chemistry Biol Chem 257: 5560–5567

Painter RC, Roseboom TJ & Bleker OP (2005) Prenatal exposure to the Dutch famine and disease in later life: An overview. Reprod Toxicol 20: 345–352

Pembrey ME, Bygren LO, Kaati G, Edvinsson S, Northstone K, Sjöström M & Golding J (2006) Sex-specific, male-line transgenerational responses in humans. Euro J Hum Gen 14: 159–166

Pointud JC, Mengus G, Brancorsini S, Monaco L, Parvinen M, Sassone-Corsi P & Davidson I (2003) The intracellular localisation of TAF7L, a paralogue of transcription factor TFIID subunit TAF7, is developmentally regulated during male germ-cell differentiation. J Cell Sci 116: 1847–1858

Prados J, Stenz L, Somm E, Stouder C, Dayer A & Paoloni-Giacobino A (2015) Prenatal exposure to DEHP affects spermatogenesis and sperm DNA methylation in a strain-dependent manner. PLoS ONE 10: e0132136

Quinnies KM, Doyle TJ, Kim KH & Rissman EF (2015) Transgenerational effects of Di-(2-Ethylhexyl) phthalate (DEHP) on stress hormones and behavior. Endocrinology 156: 3077–3083

Radford EJ, Ito M, Shi H, Corish JA, Yamazawa K, Isganaitis E, Seisenberger S, Hore TA, Reik W, Erkek S, et al (2014) In utero undernourishment perturbs the adult sperm methylome and intergenerational metabolism. Science 345: 1255903

Rechavi O, Houri-Ze’Evi L, Anava S, Goh WSS, Kerk SY, Hannon GJ & Hobert O (2014) Starvation-induced transgenerational inheritance of small RNAs in C. elegans. Cell 158: 277–287

Reddi PP, Naaby-Hansen S, Aguolnik I, Tsai JY, Silver LM, Flickinger CJ & Herret JC (1995) Complementary deoxyribonucleic acid cloning and characterization of mSP-10: the mouse homologue of human acrosomal protein SP-10. Biol Reprod 53: 873–81

Seong KH, Li D, Shimizu H, Nakamura R & Ishii S (2011) Inheritance of stress-induced, ATF-2-dependent epigenetic change. Cell 145: 1049–1061

Shinagawa T, Huynh LM, Takagi T, Tsukamoto D, Tomaru C, Kwak HG, Dohmae N, Noguchi J & Ishii S (2015) Disruption of TH2a and TH2b genes causes defects in spermatogenesis. Development 142: 1287–1292

Skinner MK, Nilsson E, Sadler-Riggleman I, Beck D, ben Maamar M & McCarrey JR (2019) Transgenerational sperm DNA methylation epimutation developmental origins following ancestral vinclozolin exposure. Epigenetics 14: 721–739

Song Y, Wu N, Wang S, Gao M, Song P, Lou J, Tan Y & Liu K (2014) Transgenerational impaired male fertility with an Igf2 epigenetic defect in the rat are induced by the endocrine disruptor p,p’-DDE. Hum Reprod 29:2512–2521

Stenz L, Escoffier J, Rahban R, Nef S & Paoloni-Giacobino A (2017) Testicular dysgenesis syndrome and Long-Lasting epigenetic silencing of mouse sperm genes involved in the reproductive system after prenatal exposure to DEHP. PLoS ONE 12: e0170441

Stenzid L, Rahban R, Pradosid J, Nef S & Giacobino AP (2019) Genetic resistance to dehp-induced transgenerational endocrine disruption. PLoS ONE 14: e0208371

Taouk L & Schulkin J (2016) Transgenerational transmission of pregestational and prenatal experience: Maternal adversity, enrichment, and underlying epigenetic and environmental mechanisms. J DevOrig Health Dis 7: 588–601

Tobi EW, Lumey LH, Talens RP, Kremer D, Putter H, Stein AD, Slagboom PE & Heijmans BT (2009) DNA methylation differences after exposure to prenatal famine are common and timing- and sex-specific. Hum Molr Gens 18: 4046–4053

Ungewitter E, Rotgers E, Bantukul T, Kawakami Y, Kissling GE & Yao HHC (2017) Teratogenic effects of in utero exposure to Di-(2-Ethylhexyl)-Phthalate (DEHP) in B6:129S4 Mice. Toxicol Sci 157: 8–19

de Vries FAT, de Boer E, van den Bosch M, Baarends WM, Ooms M, Yuan L, Liu JG, van Zeeland AA, Heyting C & Pastink A (2005) Mouse Sycp1 functions in synaptonemal complex assembly, meiotic recombination, and XY body formation. Genes Dev 19: 1376–1389

Waterland RA & Jirtle RL (2003) Transposable Elements: Targets for Early Nutritional Effects on Epigenetic Gene Regulation. Mol Cell Biol 23: 5293–5300

Wei Y, Yang CR, Wei YP, Zhao ZA, Hou Y, Schatten H & Sun QY (2014) Paternally induced transgenerational inheritance of susceptibility to diabetes in mammals. Proc Natl Acad Sci U S A 111: 1873–1878

Wen Y, Rattan S, Flaws JA & Irudayaraj J (2020) Multi and transgenerational epigenetic effects of di-(2-ethylhexyl) phthalate (DEHP) in liver. Toxicology and Applied Pharmacology 402: 115123

Xia L, Hou Y, Li Y & Cao D (2021) Transgenerational male reproductive effect of prenatal arsenic exposure: abnormal spermatogenesis with Igf2/H19 epigenetic alteration in CD1 mouse. Int J Environ Health Res 1–13 Online ahead of print

Yamaguchi K, Hada M, Fukuda Y, Inoue E, Makino Y, Katou Y, Shirahige K & Okada Y (2018) Re-evaluating the Localization of Sperm-Retained Histones Revealed the Modification-Dependent Accumulation in Specific Genome Regions. Cell Reports 23: 3920–3932

Yoshida K, Maekawa T, Ly NH, Fujita S ichiro, Muratani M, Ando M, Katou Y, Araki H, Miura F, Shirahige K, et al (2020) ATF7-Dependent Epigenetic Changes Are Required for the Intergenerational Effect of a Paternal Low-Protein Diet. Mol Cell 78: 445–458.e6

Yoshimizu T, Sugiyama N, de Felice M, Yeom Y, Ohbo K, Masuko K, Obinata M, Kuniya A, Schöler HR & Matsui Y (1999) Germline-specific expression of the Oct-4/green fluorescent protein (GFP) transgene in mice. Dev Growth Diff 41: 675–684

Zeybel M, Hardy T, Wong YK, Mathers JC, Fox CR, Gackowska A, Oakley F, Burt AD, Wilson CL, Anstee QM, et al (2012) Multigenerational epigenetic adaptation of the hepatic wound-healing response. Nat Med 18:1369–1377

